# Machine learning algorithms predict soil seed bank persistence from easily available traits

**DOI:** 10.1101/2022.01.19.476872

**Authors:** Sergey Rosbakh, Maximilian Pichler, Peter Poschlod

**Affiliations:** Ecology and Nature Conservation Biology, University of Regensburg, Germany; Theoretical Ecology, University of Regensburg, Germany

**Keywords:** artificial intelligence, predictive modeling, persistence, random forest, seed, soil, trait

## Abstract

**Question:** Soil seed banks (SSB), i.e., pools of viable seeds in the soil and its surface, play a crucial role in plant biology and ecology. Information on seed persistence in soil is of great importance for fundamental and applied research, yet compiling datasets on this trait still requires enormous efforts. We asked whether the machine learning (ML) approach could be used to infer and predict SSB properties of a regional flora based on easily available data.

**Location:** Eighteen calcareous grasslands located along an elevational gradient of almost 2000 m in the Bavarian Alps, Germany.

**Methods:** We compared a commonly used ML model (random forest) with a conventional model (linear regression model) as to their ability to predict SSB presence/absence and density using empirical data on SSB characteristics (environmental, seed traits and phylogenetic predictors). Further, we identified the most important determinants of seed persistence in soil for predicting qualitative and quantitative SSB characteristics using the ML approach.

**Results:** We demonstrated that the ML model predicts SSB characteristics significantly better than the linear regression model. A single set of predictors (either environment, or seed traits, or phylogenetic eigenvectors) was sufficient for the ML model to achieve high performance in predicting SSB characteristics. Importantly, we established that a few widely available SSB predictors can achieve high predictive power in the ML approach, suggesting a high flexibility of the developed approach for use in various study systems.

**Conclusions:** Our study provides a novel methodological approach that combines empirical knowledge on the determinants of SSB characteristics with a modern, flexible statistical approach based on ML. It clearly demonstrates that ML can be developed into a key tool to facilitate labor-intensive, costly and time-consuming functional trait research.

## Introduction

Soil seed banks (SSB), pools of viable seeds in the soil and its surface, play a key role in plant biology and ecology at different levels of organization. They bridge short- and long-term environmental conditions temporarily unsuitable for growth and reproduction, especially in habitats subject to high climatic variability and high levels of disturbance, competition and predation (Fenner et al., 2005; Saatkamp et al., 2014). Regeneration resulting from persistent SSBs helps plants to recover the original state of populations and communities, including genetic diversity (Honnay et al., 2008), after they have been altered by environmental fluctuations (Vandvik et al., 2016). Thus, the ability of seeds to persist in the soil for long periods is a crucial bet-hedging strategy (Harper, 1977; Venable & Brown, 1988; Rosbakh & Poschlod, 2021) contributing to a plant’s adaptive potential and ecosystem resilience, which is especially important in times of global change (Walck et al., 2011; Ooi, 2012).

Species that are not able to persist in the soil at a local or regional scale are particularly vulnerable to extinction risk, whereas species with persistent SSBs can easily recover even after direct destruction of above-ground vegetation (Stöcklin & Fischer, 1999; Plue & Cousins, 2018; Plue et al., 2020). Thus, knowledge about species’ ability to form persistent SSBs is of great importance for fundamental and applied research, such as nature conservation or restoration (Bakker et al., 1996; Willems & Bik, 1998; Faist et al., 2013) and invasive species management (Gioria et al., 2019). However, compiling databases on seed persistence in soil requires enormous effort when collecting primary data: such studies are labor-intensive, costly, and time-consuming. As a result, the existing studies are limited to a few regions (e.g., temperate Europe; Plue et al., 2020) and specific habitats (e.g., grasslands; Kleyer et al. 2018). Consequently, life-history trait databases suffer from a chronic problem of missing data on seed persistence in soil. We are simply uncertain about the survival potential of species in the soil in entire local and regional floras, which impedes, for example, extinction risk assessment studies (Stöcklin & Fischer, 1999), research on community assembly (Jiménez-Alfaro et al., 2016), and habitat restoration programs (Hölzel & Otte, 2004).

Several characteristics determine seed persistence in the soil, including seed and whole plant traits, vegetation and environmental properties, and various combinations thereof (Poschlod et al., 2013; Saatkamp et al., 2014). To begin with, morphological (seed shape and seed size) and physiological traits (dormancy) have been widely used as predictors of seed persistence in soil: species with comparatively small, round, dormant seeds tend to build persistent and dense(r) banks in the soil (Bekker et al., 1998; Honda, 2008; Gioria et al., 2020). Further, seed number (i.e., total seed production per individual plant), species population density and its dominance in the vegetation are considered important, especially for species that form SSBs, as species that produce a high number of seeds (often annuals; Phartyal, et al., 2020) and/or dominate in the vegetation canopy tend to have denser SSBs (Arroyo et al., 1999; Hölzel & Otte, 2004; Gioria et al., 2019).

Importantly, the predictive power of these characteristics varies strongly with local environmental conditions, suggesting that abiotic and biotic factors can mediate species ability to form SSBs (Abedi et al., 2014; Saatkamp et al., 2014; Long et al., 2015; Rosbakh & Poschlod, 2021;). In general, species richness, composition, and density of the SSBs are positively correlated with conditions of unpredictable growth, frequent disturbance and high-risk recruitment (Anderson et al., 2012; Gioria et al., 2020). Previous research on seed bank variation across successional gradients in different habitats indicates that the persistence and size of SSBs decrease with successional maturity (Warr et al., 1994; Gioria et al., 2020; Plue et al., 2020). Additionally, a few existing studies on SSB variability along environmental gradients have revealed that all characteristics of SSBs, but particularly seed density, are negatively correlated with levels of abiotic stress, e.g., climate and edaphic conditions (Funes et al., 2003), due to their direct and indirect effects on seed persistence in the soil (Fenner et al., 2005; Poschlod et al., 2013; Saatkamp et al., 2014; Long et al., 2015). Finally, the recent study by Gioria et al. (2020) demonstrated that SSB type and density depends on species relatedness, suggesting that the ability to form persistent and/or dense SSBs might be inferred from phylogeny.

Such interconnected and variable relationships between predictors of SSB persistence (plant and seed traits, habitat preferences and phylogeny) make predicting SSB characteristics challenging. In particular, conventional statistical methods, such as regression models, are not suitable for this task as their learning, i.e., finding a relationship between the predictors (e.g., seed traits) and the response (SSB characteristics), is guided and constrained by *a priori* assumption(s) about the underlying relationships, thereby limiting their predictions with a pre-defined set of rules, which for SSB characteristics are currently only poorly understood. In this context, Machine Learning (ML) is a promising tool for solving this problem.

Modern ML algorithms can flexibly identify the best predictive predictors, non-linearities of predictors, and interactions between predictors, and usually achieve higher predictive performance than regression models (Breiman, 2001b; LeCun, Bengio, & Hinton, 2015). Recent studies have demonstrated that ML models can successfully predict plant-environment relationships and outperform conventional methods, for example GLMs (Pichler et al., 2020), by a substantial margin. Moreover, ML models cope well with high-dimensional data. And yet, the ML approach has never been applied to infer and predict species persistence in SSBs. Finally, many potential predictors of SSB persistence have been identified in recent years, but it remains unclear which characteristics contribute the most to predicting SSB persistence. This is an important question since collecting data on seed and plant traits, phylogeny, and environmental data results in different costs. ML could offer an attractive solution, as most likely multiple predictors that exist for SSB are difficult to identify with regression models, while ML is more flexible and efficient in detecting the most predictive patterns.

The main aim of this study is to test the applicability of the ML approach to infer and predict SSB properties in a regional flora. Specifically, we ask two questions: 1) Can we predict species’ abilities to build a SSB (and its density) better with ML than with commonly used (generalized) linear models? 2) What determinants of seed persistence in soil (environmental characteristics, seed traits and phylogenetic relatedness) are the most important for predicting qualitative and quantitative SSB characteristics using the ML approach? The practical utility of this approach in seed ecological research is demonstrated using an extensive SSB survey of a set of easily available seed and plant traits, and species phylogeny and environmental characteristics conducted in 18 species-rich grasslands located along a climatic gradient. The study is also intended to provide detailed explanations of the methods used in order to stimulate further usage of the ML approach in seed science research.

## Materials and methods

### Study system

The field data were collected from species-rich calcareous grasslands on nutrient-poor soil located along an elevational gradient in the Bavarian Alps (northern part of the Calcareous Alps, Germany; Appendix S1) from 656 to 2363 meters above sea level. We selected this study system due to two main reasons. First, these ecosystems are ideal for studying the relative impacts of environmental (un)favorability on SSBs because elevation gradient encompasses strong variation in climatic factors (temperature), soil conditions (soil moisture and nutrients), disturbance regimes (substrate stability, past and present land-use type) and many other environmental properties (Körner, 2007) potentially affecting seed persistence in soil. Second, the relatively high number of taxonomically and functionally diverse species occurring in the studied grasslands allowed us to test the influence of plant phylogeny and seed traits on seed persistence in soil.

The study region is typical for the Northern Alps in Southern Germany, with steep Triassic lime and dolomite mountain peaks. The climate has mean annual precipitation rates up to 1500-2000 mm/year and a strong altitudinal decrease in mean annual temperature of ca. -0.6°C/100 m of elevation (Marke et al., 2013). The lower montane vegetation is dominated by tall forbs and grasses, which are replaced by sedges, short-stature herbs and dwarf shrubs as altitude increases. During the first half of the 20^th^ century, the traditional practice of grazing and mowing ceased, although several study sites were occasionally grazed by cattle or wild ungulates. The nomenclature follows Oberdorfer (2001).

### Soil seed bank survey

In 2009, we selected 18 sites (Appendix S2) located at different elevations representing different grassland vegetation types typical for the study region and easily accessible by foot for soil sample transportation. The SSBs were studied by cultivating the soil samples in an open greenhouse in Regensburg. More specifically, the soil samples were collected right after snowmelt: from the beginning of April to the second half of May in the years 2010 – 2017 (the sampling period is elevation specific). The sampling period was spread over eight years due to limited space for soil sample cultivation. We assumed that the studied SSBs are rarely subject to considerable year-to-year fluctuations, as the disturbance levels in the study system are very low and succession rates are slow. Thus it is most likely that sampling over different years did not affect the SSB characteristics. At each site, we randomly selected ten 2m x 2m plots (replicates) with homogenous vegetation. The plots were located within more or less similar distances from each other within an area of ca. 1000 m² at each site. At each plot, soil was cored with a soil auger (4 cm diameter) to a maximum depth at 10 random locations and the samples were bulked together. The top layer of each soil core including the litter layer and the top centimeter were removed to exclude transient seeds present at the surface. We focused on the top 10 cm of the soil profile to account for elevation-specific differences in the sampled volume of soil, as lowland grasslands tend to have deeper soils as compared to their upland counterparts. A preliminary study conducted in a few lowland sites indicated that this approach would not affect the correctness of the SSB characteristics, as very few viable seeds were found below the first 10 cm of the soil profile (S. Rosbakh, unpublished data). Altogether, there were 100 soil samples from each site, resulting in 1800 samples in total.

The collected soil samples were transported to the lab, where they were stored at +4 °С for a few days before being processed. The soil samples were bulked by sieving through a 0.2-mm sieve, spread thinly and evenly on plastic trays (40 cm wide) filled with potting soil, and cultivated outdoors at the University of Regensburg (Germany). To allow all viable seeds to germinate, the samples were cultivated for two successive growing seasons. Emerged seedlings were identified and removed from the trays. Five containers with potting soil only were used to control for contamination by airborne seeds or seeds present in the potting soil. After the cessation of the initial flush of germination during the first cultivation year, the soil samples were carefully turned over with a fork to facilitate the germination of ungerminated seeds. After cold stratification during the winter between two growing seasons, the soil samples were turned over one more time. Cultivation was discontinued when no more seedlings emerged for eight consecutive weeks.

### SSB predictors

#### Environmental characteristics of the study sites

We considered three main types of SSB predictors: environmental factors, seed traits and phylogeny. Environmental predictors included thermal conditions, water and nutrient supply, and disturbance (grazing). Abundances of individual species in the vegetation at each site were included in the group of environmental predictors as they can be considered to be a result of abiotic filtering. The vegetation was surveyed in the same plots from which the soil samples were taken. The surveys were conducted in the same year that the soil was sampled in ten 2 x 2 plots per site at the peak of the growing season, which was elevation specific. In each plot, the abundance of all vascular plant species was estimated based on the following scale: 0.1-1, 1-5, 5-28, 25-50, 50-75, and 75-100%. The relative abundance of a species at a site was then calculated as the mean value of its abundance in all plots.

Site thermal conditions during the vegetation period were estimated with the help of the Landolt indicator value for temperature (Landolt’s T), a proxy for mean soil and surface temperatures after snow-melt (Landolt et al., 2010; Scherrer & Körner, 2011). Similarly, we used Landolt indicator values for water availability (Landolt’s F) and soil nutrients (Landolt’s N) as proxies for site water and nutrient supply during the vegetation period, respectively. We opted for these indicator values because they are strongly correlated with directly measured temperature and soil parameters (e.g., air temperature, soil phosphorous content, soil depth; e.g., Rosbakh and Poschlod 2021) and due to their wider availability. Finally, grazing intensity, the main disturbance factor at the study sites, was recorded at all study sites and included three levels: 1) no current agricultural usage but occasional grazing by sheep and wild ungulates, 2) occasional extensive grazing by cows, and 3) mountain dairy farm with permanently grazing cows (except for HO5 site extensively grazed by sheep).

### Seed traits

For all the species found either in the SSBs or in the above-ground vegetation we collected *in situ* data on seed mass, seed shape and seed production following the standardized protocols (Kleyer et al., 2008). Seed mass was extrapolated based on three samples of 100 seeds each.

Seed shape is the variance of seed dimensions that were measured on ten replicate seeds per species. It is a dimensionless trait that varies between zero in perfectly round and 0.2 in disk- or needle-shaped seeds. Seed production was measured as the average number of seeds produced per ramet of ten randomly selected individuals (Thompson et al., 1993).

Furthermore, all the species were classified into having either non-dormant or dormant seeds based on published literature (Baskin & Baskin, 2014; Rosbakh et al., 2020) and own germination tests (S. Rosbakh, unpublished). Finally, the presence/absence of endosperm was identified for every species according to (Martin, 1946; Finch-Savage & Leubner-Metzger, 2006;).

Trait data on seed mass, seed shape and seed production were unavailable for some (less than 15) of the species; in their case the missing data were extracted from the LEDA database (Kleyer et al., 2008).

### Phylogeny

To infer the influence of species’ phylogenetic relatedness on seed persistence in soil, e.g., Gioria et al. (2020), we included information on the phylogenetic distances between study species as variables in the models. We made no inferences about the potential evolutionary processes underlying possible correlation between SSB properties and species phylogeny. The phylogenetic relationships among all the studied species were summarized by calculating eigenvectors extracted from a principal coordinate analysis (PCoA), representing the variation in the phylogenetic distances among species (Penone et al., 2014). We used the first 13 eigenvectors that represented more than 60% of the variation in the phylogenetic distances among species. The calculation of the eigenvectors was based on a dated phylogeny of a large European flora (Durka & Michalski, 2012).

### Data analysis

All statistical calculations were done with the help of R software (version 4.1, R Core Development Team, 2019).

### Data preparation

Based on the vegetation survey and soil cultivation data, we predicted the ability of a species to form a persistent SSB at a study site as a binary variable (1 – able to form a seed bank, 0 – otherwise). Furthermore, we predicted SSB density (seeds/m2), a quantitative measure of persistence in soil, for each species at each study site by adding up the numbers of seedlings germinated from the corresponding soil samples.

In the first step, we compiled a data set including soil seed bank data (both the binary variable for ability to build a seed bank and seed bank density), environmental characteristics, seed traits, and phylogenetic relatedness for each species occurring at each study site both in the vegetation and in the soil seed bank. In other words, the analyzed dataset contained seed bank data for multiple species at the same plot, i.e., every row in the dataset represented a species-plot combination.

We transformed the ‘seed dormancy’ ordinal variable into a continuous variable. Missing values in the dataset (seed shape for 5 species, productivity for 45 species and dormancy for 37 species) were imputed using the *missRanger* R package (an alternative implementation of the original proposed method (Stekhoven & Bühlmann, 2011)). As information about vegetation succession is rarely available in soil seed bank research (many species from previous succession stages can survive in the soil for longer periods of time), the observations with vegetation equal to zero were removed from the dataset. All predictors were standardized (centered and divided by their standard deviation) prior to analysis. Because SSB density was heavily skewed, we applied logarithmic transformation to it (*log(SSBdensity* + 0.001)) and used the log-transformed variable as a response variable in our models. Model assumptions were met in all cases, when applicable.

### Model evaluation

Evaluating models on the data on which they have been trained leads to underestimation of the actual predictive error for new data (Roberts et al., 2017). To estimate the generalization ability of a model (i.e., how accurate the predictions of a model are for new observations), it has to be evaluated on a part of the dataset that was not used for training the model, the so-called holdout. We used k-folded cross-validation (i.e., split the dataset into several holdouts so that each data point appears once in the holdout dataset, trained the model *n* times on the *n* training datasets, and averaged the predictive errors on the *n* holdout datasets, see Roberts et al. (2017)).

While cross-validation can produce accurate estimates of predictive performance, performance can still be overestimated if the observations are non-independent, for example in the presence of spatial auto-correlation (Roberts et al., 2017). To counteract this, we used 9-folded, spatially blocked cross-validation to account for spatial dependencies introduced by the 18 sites from which the observations were collected. In each split, observations for 16 sites were used to train the model and the holdouts of two sites were used to estimate the predictive error. We used 9-folded blocked cross-validation for all the different sets of predictors.

For the calculation of the predictive error/performance (on the holdouts of the cross-validation), we used the area under the ROC curve (AUC) for models when predicting the presence/absence of SSB, and the R-squared for models when predicting SSB density. AUC measures how well the model can differentiate between two response classes (presence and absence of SSB). The AUC and R-squared were averaged over the 9 holdouts of the cross-validation.

### Performance of the ML and the conventional approach in predicting SSB characteristics

To test whether the ML approach is more advantageous than conventional approaches to predicting SSB characteristics, we used two common representatives of these groups. For ML, we used the random forest model (RF; Breiman, 2001a) which has advantages over other ML models such as the low number of hyper-parameters and the associated easier usability. Hyper-parameters are parameters of the model itself (not to be confused with parameters that are optimized by the model), which are usually optimized in a trial-and-error search to find the optimal set for a specific dataset (Claesen & Moor, 2015). In addition, RF copes well with small datasets, can handle different types of responses (e.g., presence/absence of SSB and SSB density in our study), and is implemented in numerous programming languages. Foregoing the established procedures, we skipped hyper-parameter optimization, opting instead to test the achievable predictive performance with the default hyper-parameters because hyper-parameter tuning usually requires expert knowledge. We used the RF implementation from the *ranger* R package (version 0.12.1 Wright, Wager, & Probst, 2016).

For the conventional statistical approach, we used linear regression (with log-transformed SSB density as the response variable) and logistic regression models (presence/absence of SB as the response variable) as these are commonly used tools in analyses of ecological data. The training or ‘learning’ in regression models is specified by the hypothesis. Because linear regression models cannot learn outside of their hypothesis, i.e., if interactions are not specified the model cannot account for them, we added all the predictors additively, as well as all the combinations of predictor-predictor interactions. To compensate for the lack of power (interim results showed that the regression model would not converge with so many predictors), we applied elastic-net regularization (Zou and Hastie, 2005) via the *glmnet* R package (Friedman et al., 2010). The strength of the regularization and the weighting between the l1 and the l2 regularization were tuned via three-fold cross-validation.

We used the *mlr* R package (version 0.9.0, (Lang et al., 2019)) to train and evaluate the models.

### Relative importance of environmental characteristics, seed traits and phylogeny in predicting SSB characteristics

We identified the relative importance of single predictors and corresponding functional groups (seed traits, environment and phylogeny) using the RF models as they demonstrated better performance than the regression framework (see below).

### Identifying individual important predictors

RF provides quantitative information about the importance of the predictors. This ranking, called variable importance (Breiman, 2001a), should be not confused with regression coefficients in regression models, since the absolute values of those variables’ importance are uninformative and depend on the dataset (and the number of predictors). However, the relative importance of the variables versus each other can be used to rank the predictors to identify the most predictive ones. Thus, to identify the most important predictors, we fitted RF on all the predictors and ranked the importance of the predictors based on their variable importance.

To assess the ability of the different functional groups to predict SSB, we divided the predictors into ‘Environment’ (Landolt’s T, Landolt’s F, Landolt’s N, grazing intensity, cover), ‘Seed’ (mass, shape, production, dormancy, endosperm presence/absence), ‘Phylogeny’ (the first 13 phylogenetic eigenvectors), and ‘All’ (all predictors). We then fitted the models (RF and regression) on the different groups and estimated the predictive performance via 9-folded blocked cross-validation (see above).

### Minimal requirements for predicting SSB features with the ML approach

The choice of type and number of model predictors in ecological research strongly depends on the available data. Thus, to estimate the minimal set of SSB determinants required to achieve high predictive performance for SSB characteristics, we selected the four previously identified predictors (see corresponding sections) with the highest variable importance, which were [temperature, c5, c7, mass] for the presence/absence of SSB and [temperature, c7, mass, c5] for SSB density, to test their predictive performance.

In the second step, we first tested an RF model with only the first predictor (temperature) and in subsequent steps we sequentially added the rest of the four predictors to the set of predictors. In each step, we estimated the predictive performance via 9-folded blocked cross-validation as described above.

### Functional relationship of important predictors and SSBs

ML models are often referred to as black-box models because it remains unknown what relationships the ML model learns in order to generate predictions. In linear regression models, the *a priori* hypothesis restricts the model’s learning, and the model is not capable of learning outside of this hypothesis (e.g., given two predictors A and B, if the interaction of A and B is not specified, the model cannot learn it). In ML, however, the idea is that the model should be capable of automatically identifying the best predictive patterns in the data (Breiman, 2001a), which makes ML a great tool for predictive modeling but comes with the cost of low interpretability (Breiman, 2001b). However, findings of discriminative ML models have driven the development of explainable AI (xAI) methods and tools (Barredo Arrieta et al., 2020). The idea of xAI is to reveal post-hoc the predictive patterns used by the ML model (Barredo Arrieta et al., 2020; Pichler et al., 2020; Ryo et al., 2021).

To check whether the predictive patterns the ML model used to predict SSB density are ecologically plausible, we used an approach based on accumulated local effect plots (ALE; Apley & Zhu, 2020) to explore the functional relationships between predictors (temperature, shape, and mass) and the response variable (presence/absence of SSB, and density of SSB; Molnar, 2020). Briefly, ALEs are based on the idea of sampling predictors individually while keeping the other predictors fixed. If the sampled predictor is ‘important’, the predictions will be affected more strongly. Phylogenetic predictors were not considered because they cannot be linked to actual ecological mechanisms, making their interpretation pointless.

## Results

All data used in the analysis are available in Appendix S2.

### Vegetation and soil seed bank surveys

At the 18 study sites, we recorded 290 species belonging to 45 families. The most dominant families were Asteraceae (45 species), Poaceae (30 species), Cyperaceae (21 species) and Caryophyllaceae (16 species). Graminoids dominated in the vegetation of all the sites surveyed.

In total, 247 995 seedlings belonging to 162 species and 35 families germinated in the collected soil samples. Thus, germinable seeds of 128 species (e.g., *Campanula alpina*, *Ligusticum mutellina* and *Valeriana montana*) were not found in the collected soil samples. Of the species present in the SSB, seeds of 65 species, for example, *Carex flacca*, *Hypericum perforatum* and *Veronica officinalis,* were found at each site where the corresponding species occurred in the vegetation. Seeds of 97 species, such as *Alchemilla vulgaris*, *Nardus stricta*, and *Ranunculus montanus*, displayed a variable behavior in the surveyed SSBs, being present at some sites and absent from others. The seed density of species present in the SSB ranged from 8 (*Carex sylvatica*, *Potentilla aurea*) to 63603 (*Sagina saginoides*) with an average of 1617 seeds/m².

### Predictive performance of the ML and conventional approaches in predicting SSB characteristics

When comparing the performance of ML (RF) and conventional approaches (linear and generalized linear model) in predicting SSB characteristics, we found that the ML approach achieved an AUC of 86% and the GLM, an AUC of 76.8% when predicting the presence/absence of SSB (Fig. 1; intersection of the circles). In predicting the density of SSBs, the ML approach achieved an R² of 41.7%, whereas the conventional approach (linear regression model) achieved an R² of 18.9% (Fig. 1; intersection of the circles).

**Figure 1:**
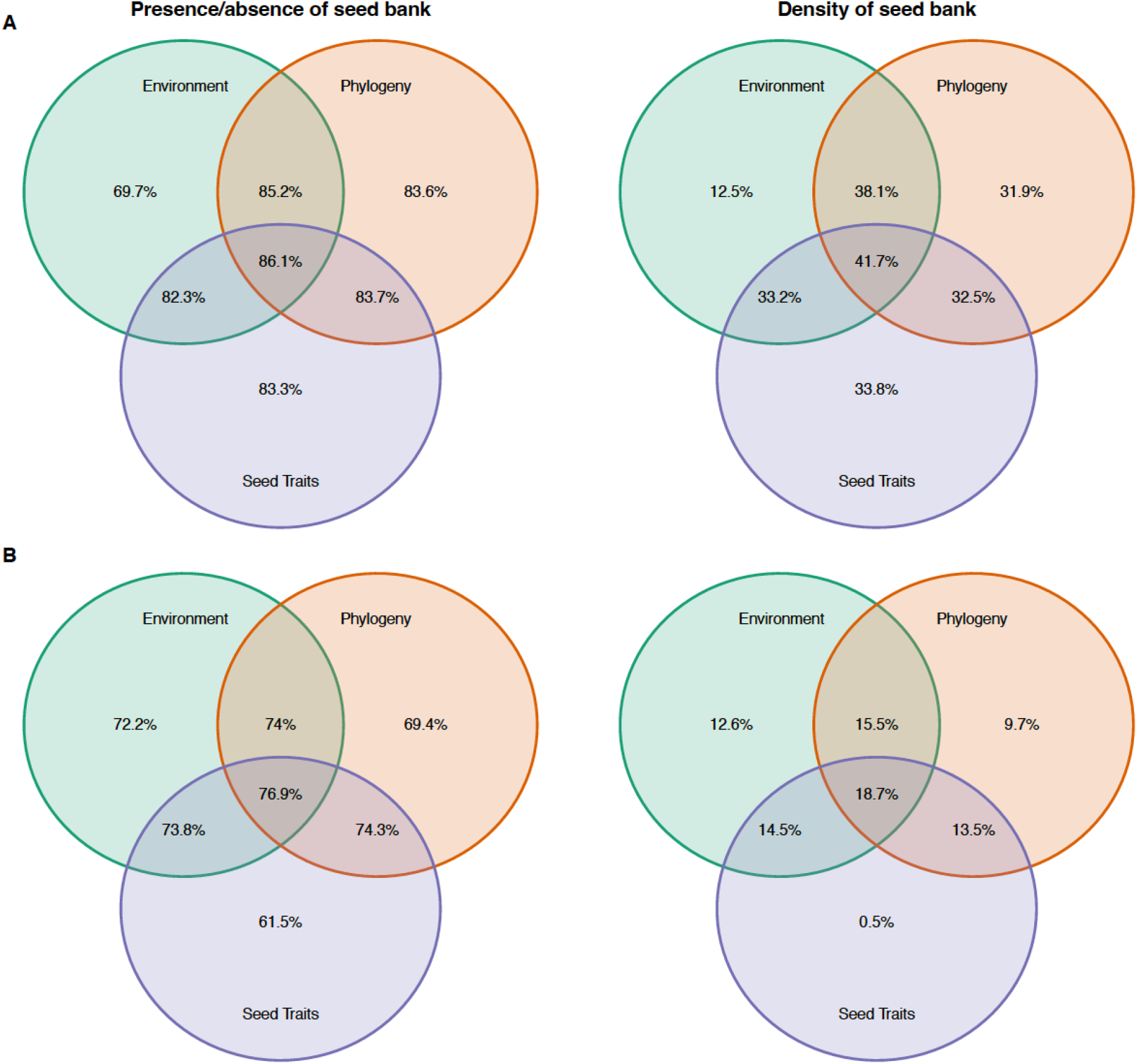
Performance of the random forest machine learning model (A) and the conventional regression model (B) in predicting presence-absence and density of seed banks. Both models were fitted on three sets of predictors (environment: temperature, nitrogen, moisture, grazing; seed traits: production, mass, endosperm, shape, and dormancy; phylogeny: phylogenetic axes which explain 60% of the variation). The intersections show the performance of the different combinations of predictors. Predictions for presence/absence of SSB (left column) were evaluated by AUC and predictions for SSB density (right column) were evaluated by R^2^. Models were evaluated by blocked nine-folded cross-validation (observations were from 18 different plots; in each validation step 16 plots were used for training and two plots for validation).

For SSB density, the combination of environmental characteristics and phylogeny included in the RF model resulted in an R^2^ of 38.6%, and was followed by the combination of environmental characteristics and seed traits (R^2^ of 32.7%), and seed traits and phylogeny (R^2^ of 32.3%). Among single groups of predictors, seed traits and phylogeny had the highest predictive performance with an R^2^ of 33.7% and 31.7%, respectively. Environmental characteristics alone were predictive of only 6.2% of SSB density in the data set.

The predictive performance of the conventional approach was substantially lower compared to the ML model, with an R^2^ of 18.9% when all predictors were used (Fig. 2B), followed by the combination of phylogenetic and seed, and phylogenetic and environmental characteristics (R^2^ of 12.7% and 16.3%). Among single groups of predictors, seed characteristics showed the lowest predictive performance with an R^2^ of 0.2%, while phylogenetic and environmental characteristics achieved higher predictive performances of 8.3% and 12.5% R^2^.

**Figure 2:**
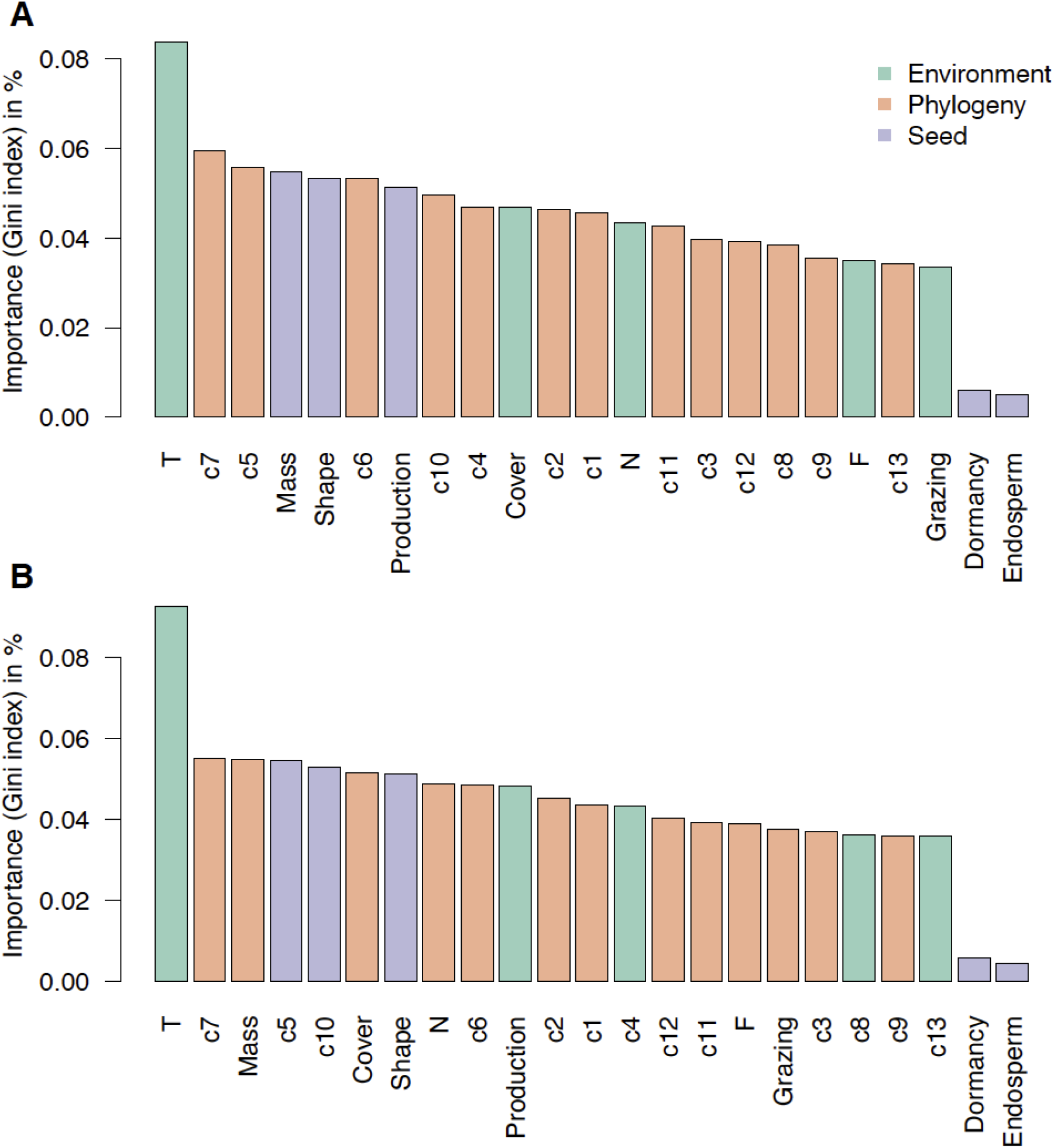
Variable importance in random forest fitted on presence/absence of seed banks (A) and on density of seed banks (B), plotted in descending order as per their relative importance measured by the Gini index in percent. All available predictors were used.

All the differences between the ML model and the (generalized) linear model were statistically significant excepting where only the group of environmental predictors was used (Appendix S3, S4, S5).

### Relative importance of environmental characteristics, seed traits and phylogeny in predicting SSB characteristics

#### Identifying individual important predictors

When looking at the variable importance of SSB predictors, we found that for both types of response (presence/absence (Fig. 2A) and density (Fig. 2B)) the temperature conditions at the surveyed sites were the most important predictor (9% and 10% for SSB presence/absence and density, respectively, Fig. 2). The remaining environmental characteristics (soil nutrients and moisture, grazing intensity), several seed traits (seed mass, shape and production) and all the phylogenetic eigenvectors had comparable variable importance for SSB characteristics (3-6%). Seed dormancy and endosperm presence/absence had comparatively low variable importance (< 1%, Fig. 2).

#### Minimal requirements for predicting SSB features with the ML approach

We identified site temperature conditions as the predictor with the highest predictive performance (AUC of 65.8% and R^2^ of 6.2%, Fig. 4) for both response types (Fig. 4). For predicting SSB presence/absence, the addition of the phylogenetic eigenvector c5 as a predictor is already sufficient to reach an AUC of 0.79, which corresponds to 91% of the maximal achievable predictive performance of 0.865 (Fig. 1A). For predicting SSB density, two additional predictors, seed mass and the phylogenetic eigenvector c5, were necessary to reach 80% of the maximal achievable predictive performance (Figs 1B and 4B).

#### Functional relationship of important predictors and SSBs

According to the RF model, the probability of a species forming an SSB increased with increasing site temperature, soil moisture and fertility, and species abundance, and decreased with increasing seed mass, seed shape value and seed production (Figs. 4A-H). Species occurring at sites with relatively high grazing intensity tended to form persistent seed banks in the soil. SSB density was positively affected by site temperature conditions, soil moisture availability and species abundance (Figs. 4I-P).

## Discussion

There is a growing demand for knowledge on soil seed persistence in both basic and applied plant ecological research. Information on species ability to form persistent SSBs and their quantitative characteristics is not only crucial for understanding past, present and future plant population dynamics (Saatkamp et al., 2014; Walck et al., 2011), but also for restoration projects (Hölzel & Otte, 2004), risk assessment (Stöcklin & Fischer, 1999) and invasive species management (Gioria et al., 2019). SSB surveys cannot by themselves satisfy such a need for knowledge as the field, and particularly the cultivation part of this approach is still extremely resource intensive.

Our study closes this gap by providing a novel methodological approach combining empirical knowledge on the determinants of SSB characteristics and a modern, flexible statistical approach based on machine learning. We first demonstrated that the ML approach substantially outperforms conventional statistical methods in predicting SSB characteristics. Second, we found that SSB characteristics can be predicted with high accuracy regardless of the available predictor type (environmental characteristics, seed trait and phylogeny). Finally, we revealed that a few widely available SSB predictors can already achieve a high predictive power in the ML approach, suggesting high flexibility of the developed approach for use in various study systems.

### Predictive performance of the ML and conventional approaches in predicting SSB characteristics

In our study, the machine learning approach (RF) outperformed the (generalized) linear model considerably in predicting SSB characteristics (Fig. 1). This finding confirms previous studies (e.g., Pichler et al. 2020) and our expectations that given the complex nature of predictors and interactions thereof, accurate analysis of patterns in SSB, and more generally ecological data requires a more flexible approach. The comparatively better performance of the ML approach in predicting SSB characteristics can be explained by two reasons.

First, assuming the correct functional form of the relationship between predictors and response variable is essential for accurate predictions. However, conventional statistical models such as linear regression models are constrained in their learning by *a priori* assumptions (the hypotheses) about the underlying system. Thus, the modeler needs to correctly specify the functional relationships between predictors and predicted characteristics (e.g., linear or non-linear) as well as the relationships between predictors. Moreover, it remains doubtful whether we can do the same for phylogenetic predictors, which can be seen as proxies for unmeasured traits (Morales-Castilla et al., 2015) and can improve the accuracy of predictions for ecological data (e.g. Brousseau et al., 2018; Pomeranz et al., 2019). However, making assumptions about their functional form or linking them to environmental or seed trait predictors is difficult as we cannot connect them to actual ecological mechanisms.

Second, conventional approaches often lack flexibility. As mentioned earlier, ecological patterns are usually scale-dependent (Poisot at al., 2015; König et al., 2021) and nuisance predictors are required to account for locally varying functional forms, but entail loss of statistical power and give no guarantee that the possible fluctuating predictive patterns of predictors will be successfully captured. However, our results demonstrated that in this case ML could offer a powerful solution, as it is able to automatically identify and learn flexible predictive patterns (Breiman, 2001a, 2001b).

Based on previous research, we assumed the existence of several predictive patterns for SSB characteristics. Our results indicate that RF individually achieved a high performance for phylogenetic and seed trait predictors (Fig. 1), but their combination did not greatly further increase predictive performance. Assuming that phylogeny is a proxy for unmeasured traits correlated with SSB persistence, our findings thus confirm that phylogeny and information about seed traits encode similar predictive patterns for SSB characteristics and using both does not increase predictive performance greatly (Fig. 1). On the other hand, for the conventional statistical models predictive performance increased greatly when all sets of functional predictors were used compared to use of individual groups (Fig. 1). This implies that such statistical models cannot make the best use of the predictive patterns in the individual groups, indicating that some of the predictive patterns are non-linear and require higher flexibility. In contrast, the ML model was able to utilize the available individual predictive patterns, highlighting the advantages of the ML approach in predicting SSB characteristics when the availability of predictors is limited by temporal and/or financial resources.

### Relative importance of environmental characteristics, seed traits and phylogeny in predicting SSB characteristics

When predicting SSB characteristics with the ML approach, we found that the temperature conditions of the surveyed sites were the most important predictor, both for SSB presence/absence and density. The conditional dependency profiles of random forest for this predictor revealed that species from warmer sites (i.e., higher Landolt’s T values) were more likely to build up a persistent SSB with higher seed density (Fig. 4). This finding is in line with our recent study in the same study system (Rosbakh & Poschlod, 2021) and observations made elsewhere (Ortega et al., 1997; Welling et al., 2004; Ma et al., 2010) that the importance of SSBs for plant persistence gradually decreases with increasing elevation. Low-temperature stress in colder sites, including the short growth period with generally low temperatures coupled with frequent and severe frost events, negatively affects regeneration by seed. Therefore, because of the unpredictable seed input into the soil, plants shift their main persistence strategy from replacement of individuals by seeding germinating from the SSB to *in situ* maintenance of established individual plants by emphasizing stasis of adult stages (Rosbakh & Poschlod, 2021).

The remaining environmental characteristics (soil nutrients and moisture, grazing intensity, and species abundance in the vegetation [‘cover’]) were found to be important predictors of SSB characteristics of equal importance, though with smaller predictive power than temperature. Although the ML approach does not allow for direct hypothesis testing and p-value calculations, which are usually used to confirm/reject postulated hypotheses, these findings are ecologically plausible as they agree well with previous SSB research. First, the detected low probability that species with persistent SSB and low SSB density would be present in sites with nutrient-poor and dry soils (i.e. lower Landolt’s F and N values) is in line with the general observation that all components of SSBs, and particularly seed density (the curves for SSB density are much steeper than for SSB presence/absence; Fig. 4), are negatively correlated with levels of abiotic stress (e.g. edaphic conditions; (Funes et al., 2003)), due to its direct and indirect effects on seed persistence in the soil (Fenner et al., 2005; Poschlod et al., 2013; Saatkamp et al., 2014; Long et al., 2015). Second, the revealed positive effects of grazing animals on SSB persistence and density agree well with the previous finding that frequent (moderate) disturbance favors formation of persistent SSBs with a high density due to the establishment of gaps by grazing and trampling, favouring species with a ruderal strategy (Grime, 2006; Renne & Tracy, 2007). Finally, SSB persistence and density were positively affected by plant abundance in the vegetation, a pattern known from other systems and explained by a comparatively large seed input into soils from the dominant species (Saatkamp et al., 2014). These results, however, come with the reservation that we did not check which predictor-predictor interactions were learned by RF. It is likely that RF found some, but the high predictive performances of a few single predictors (Fig. 4) suggest that these are negligible.

In our study, three out of five seed traits: mass, shape and production, performed well in predicting SSB characteristics. Like in other SSB studies (Honda, 2008; Bekker et al., 1998; Gioria et al., 2020), the species in our system with comparatively small, round seeds tended to build persistent and dense(r) banks in the soil, a seed morphology that favors easier seed burial and reduces risk of predation (Fenner et al., 2005). Seed production, a trait with predictive performance comparable to that of seed mass and shape, had a negative effect on SSB persistence and density, especially in species that produce more than 9000 seeds per ramet. This finding contradicts previous observations that high seed production is an important determinant of SSB characteristics due to the positive trade-off between number of produced seeds and their mass, i.e., productive species tend to produce smaller seeds that persist in the soil (Saatkamp et al., 2014).

Seed dormancy and endosperm presence played a minor role in predicting SSB characteristics, as they showed the lowest predictive performance in the calculated models. The former finding agrees well with the studies by Thompson et al., (2003) and Gioria et al. (2020), which demonstrated that seed dormancy is an important mechanism promoting seed persistence in the soil but, overall, is a poor predictor of SSB characteristics. The weak predictive power of endosperm presence in inferring SSB characteristics supports the conclusion by Long et al. (2015) that this trait, which can nevertheless serve as a good proxy for seed longevity in storage (Probert et al., 2009; Tausch et al., 2019), does not reflect species ability to persist in soil.

Including phylogenetic eigenvectors considerably improved RF model performance in predicting both SSB characteristics of interest. These results agree well with recent trait-based research showing that phylogenetic predictors contain information on unobserved traits, thereby increasing the predictive power of models (Morales-Castilla et al., 2015; Desjardins-Proulx et al., 2017; Pomeranz et al., 2019). In the SSB context, these unobserved (and usually hard-to-measure) traits might include a number of ecophysiological adaptations, such as desiccation tolerance and/or genetic degradation resistance, which positively influence inherent seed longevity and thus seed persistence in soil (Long et al., 2015). Alternatively, the good predictive performance of the phylogenetic predictors could be explained by their correlation with the seed traits correlated with SSB persistence (mass, shape productivity; Figs. 2 and 3, Appendix S6), which are not randomly distributed across phylogeny (e.g., Gioria et al. 2020). Although in our study it was not feasible to separate these two explanations from each other, we believe that in our case the latter explanation is more likely as both the ‘Seed’ and ‘Phylogeny’ groups of predictors showed the highest predictive performance of three groups but including both did not substantially improve predictive performance.

**Figure 3:**
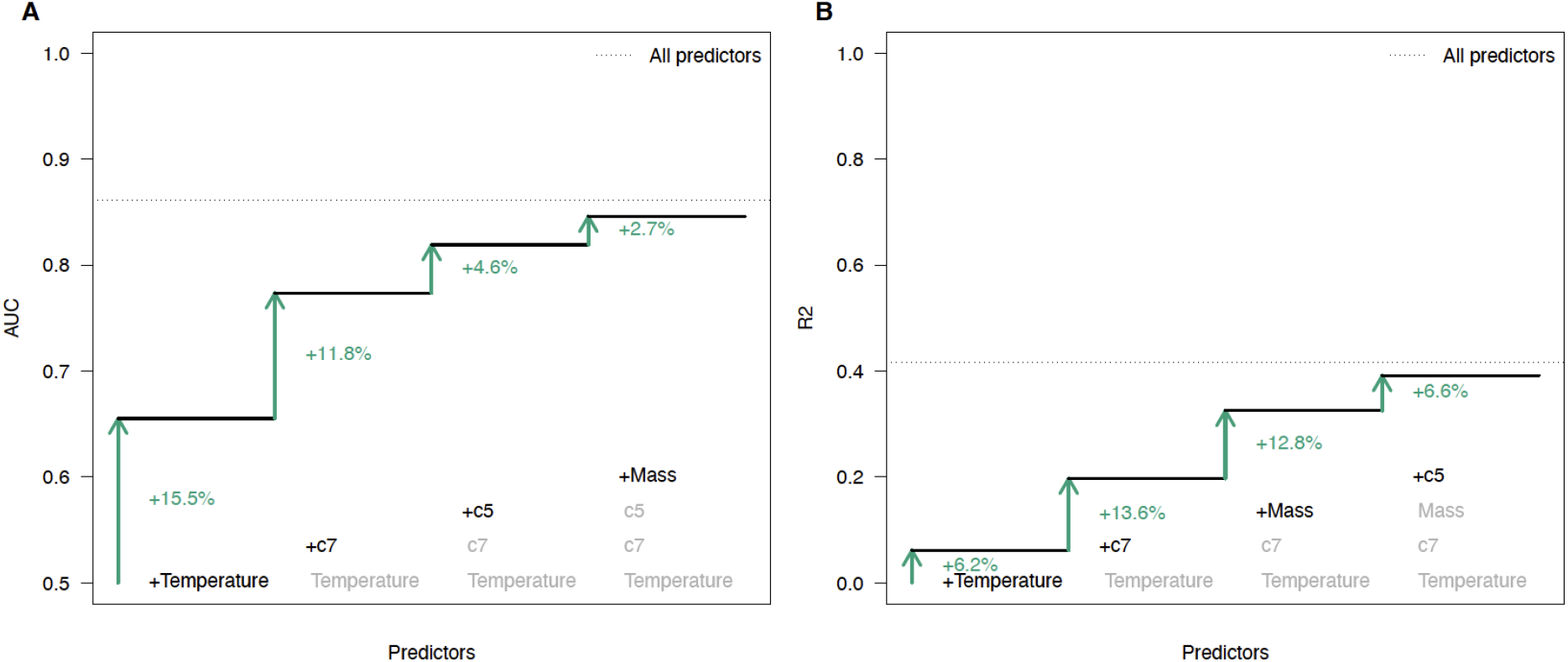
Predictive performance of random forest models for soil seed bank presence/absence (A) and density (B) with the four most important predictors.

**Figure 4:**
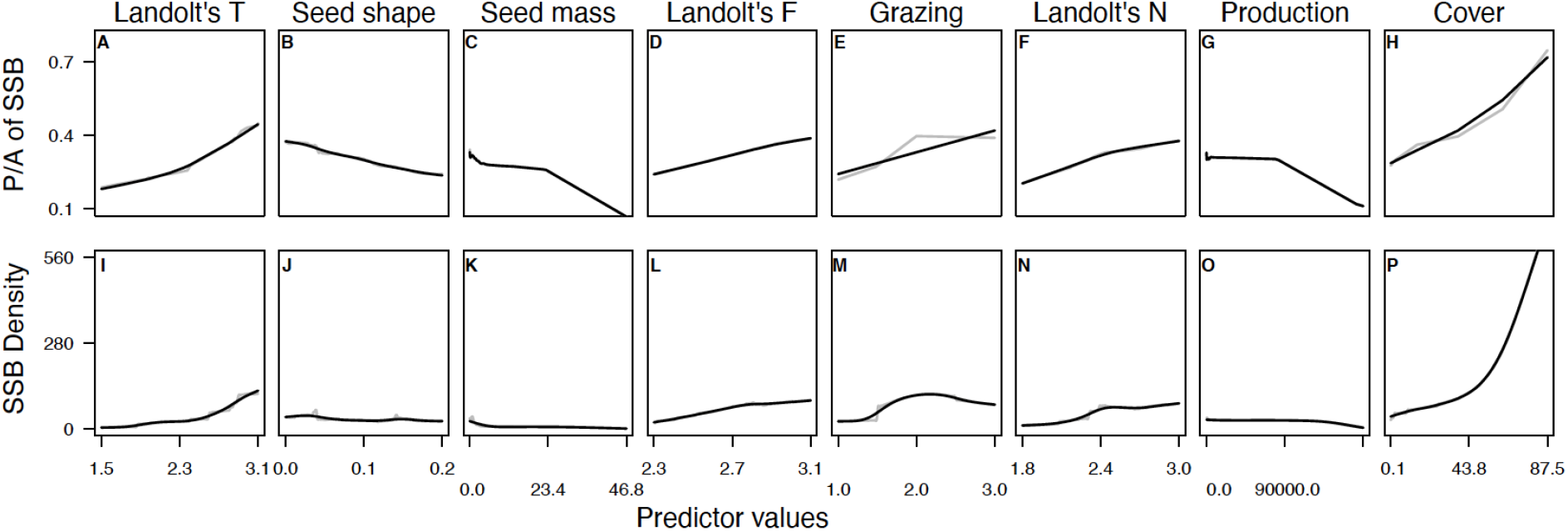
Conditional dependency profiles (based on accumulated local effects) from random forest for the environmental and the seed trait predictors. Predictors are sorted according to their variable importance found by random forest. A–H show the profiles for predicting the presence/absence of seed bank formation and I-P show the profiles for predicting seed bank density. The grey lines are the profiles, and the black lines are smoothing splines.

Besides testing different sets of predictors (environment, seed, and phylogeny), we also wanted to identify the minimal combination of the most predictive predictors independently of their group. In our study, we considered 22 predictors, a comparatively large number that would entail high labor and temporal costs of data collection, especially in poorly studied regional floras. Our results indicate that both SSB components can be predicted with high accuracy based only on a few characteristics that can be obtained from already existing sources. For example, for studies conducted in Europe, information on site temperature conditions could be obtained from regional indicator values (e.g. Landolt et al., 2010; Tyler et al., 2021), data on seed shape and mass, from trait data bases (Kleyer et al., 2008; Liu et al., 2019), and phylogenetic vectors, from the work by Durka & Michalski (2012). In other regions with poorer data coverage global ready-to-use phylogenies (e.g., Jin & Qian, 2019) in combination with *in situ* measurements of relatively simple seed morphological traits, such as mass and shape, could be used as reliable predictors of SSB characteristics.

## Authors’ contributions

SR conceived the study, MP conducted the analysis, and the first version of the manuscript was written by SR together with MP. All the authors contributed to manuscript editing. We appreciate the comments of three anonymous referees on the earlier version of the manuscript.

## Data availability

The code and data used for the analysis are publicly available at https://github.com/MaximilianPi/Rosbakh-Pichler-Poschlod-2021.

## Appendix S1.

Location of the study sites (n=18) along elevational gradient in the Bavarian Alps, Germany (see Appendix 2 for detailed environmental characteristics). Different letters indicate sampling ‘subregions’: H – ‘Hochkalter’, HO – ‘Hochkalter Ost’, M – Mordaualm, W – ‘Wimbachtal’, WM – ‘Watzmann’.

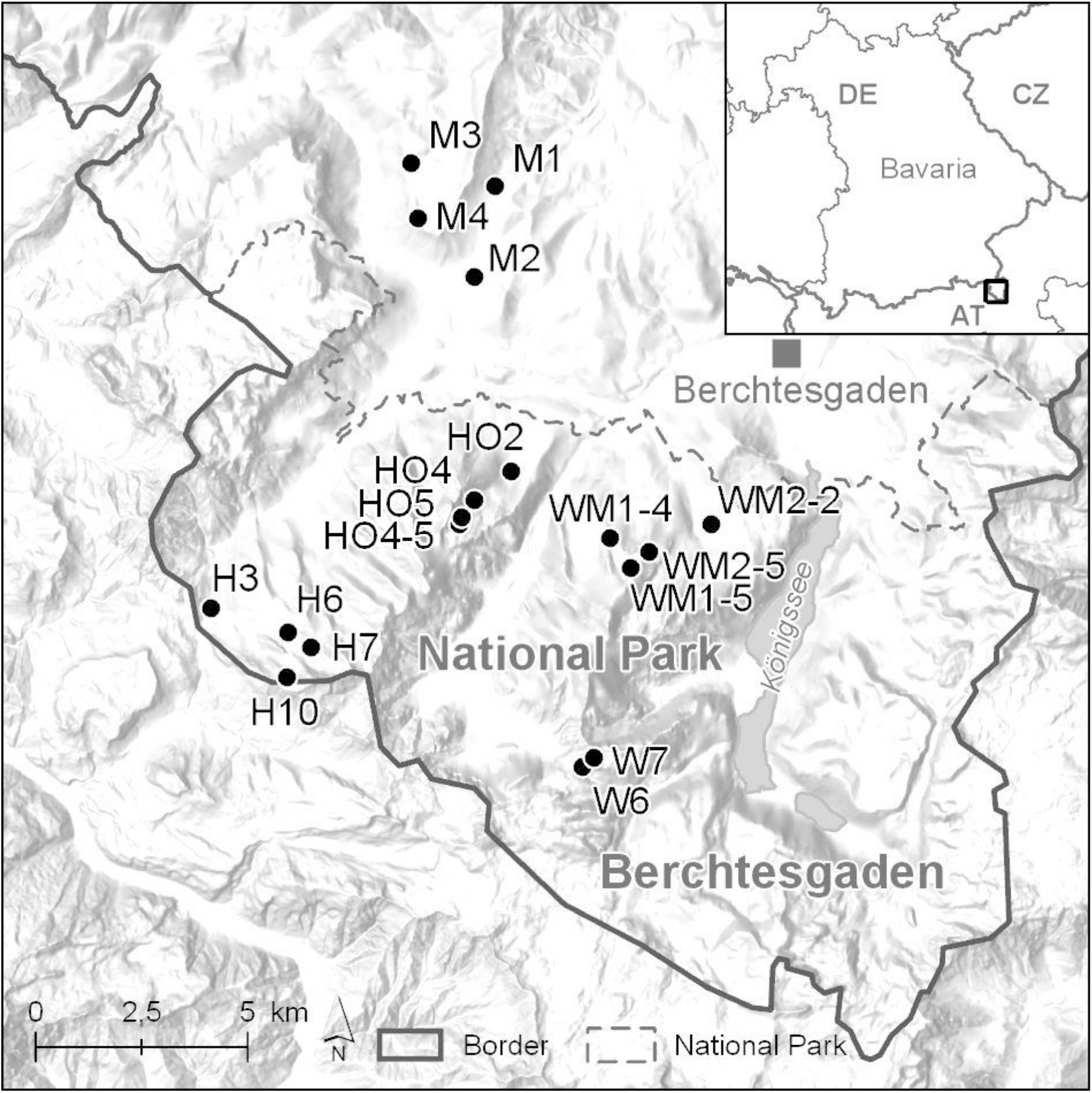

## Appendix S2.

Data used for the statistical analysis.

## Appendix S3.

Results of t-tests (p-value) comparing the random forest (RF) and generalized linear model for the different predictor groups for presence/absence of SSB.

| Scenario | lm | lm-SE | RF | RF-SE | p-value |
| --- | --- | --- | --- | --- | --- |
| All | 0.769 | 0.022 | 0.861 | 0.010 | 0.003 |
| Phylo | 0.694 | 0.019 | 0.836 | 0.013 | 0.000 |
| Seed | 0.615 | 0.015 | 0.833 | 0.010 | 0.000 |
| Env | 0.722 | 0.025 | 0.697 | 0.015 | 0.405 |
| Phylo.Seed | 0.743 | 0.026 | 0.837 | 0.012 | 0.007 |
| Phylo.Env | 0.740 | 0.021 | 0.852 | 0.012 | 0.001 |
| Env.Seed | 0.738 | 0.025 | 0.823 | 0.012 | 0.010 |

## Appendix S4.

Results of t-tests (p-value) comparing the random forest (RF) and linear regression model for the different predictor groups for SSB density.

| Scenario | lm | lm-SE | RF | RF-SE | p-value |
| --- | --- | --- | --- | --- | --- |
| All | 0.199 | 0.026 | 0.416 | 0.019 | 0.000 |
| Phylo | 0.091 | 0.022 | 0.317 | 0.026 | 0.000 |
| Seed | 0.015 | 0.007 | 0.337 | 0.023 | 0.000 |
| Env | 0.119 | 0.014 | 0.124 | 0.018 | 0.833 |
| Phylo.Seed | 0.132 | 0.026 | 0.324 | 0.028 | 0.000 |
| Phylo.Env | 0.157 | 0.022 | 0.383 | 0.022 | 0.000 |
| Env.Seed | 0.172 | 0.030 | 0.329 | 0.020 | 0.001 |

## Appendix S5.

Absolute effect estimates of generalized linear model (A) for presence/absence of SSB and of linear model (B) for SSB density. Only the 30 largest effect estimates are shown here. The elastic-net approach was applied to regularize effects.

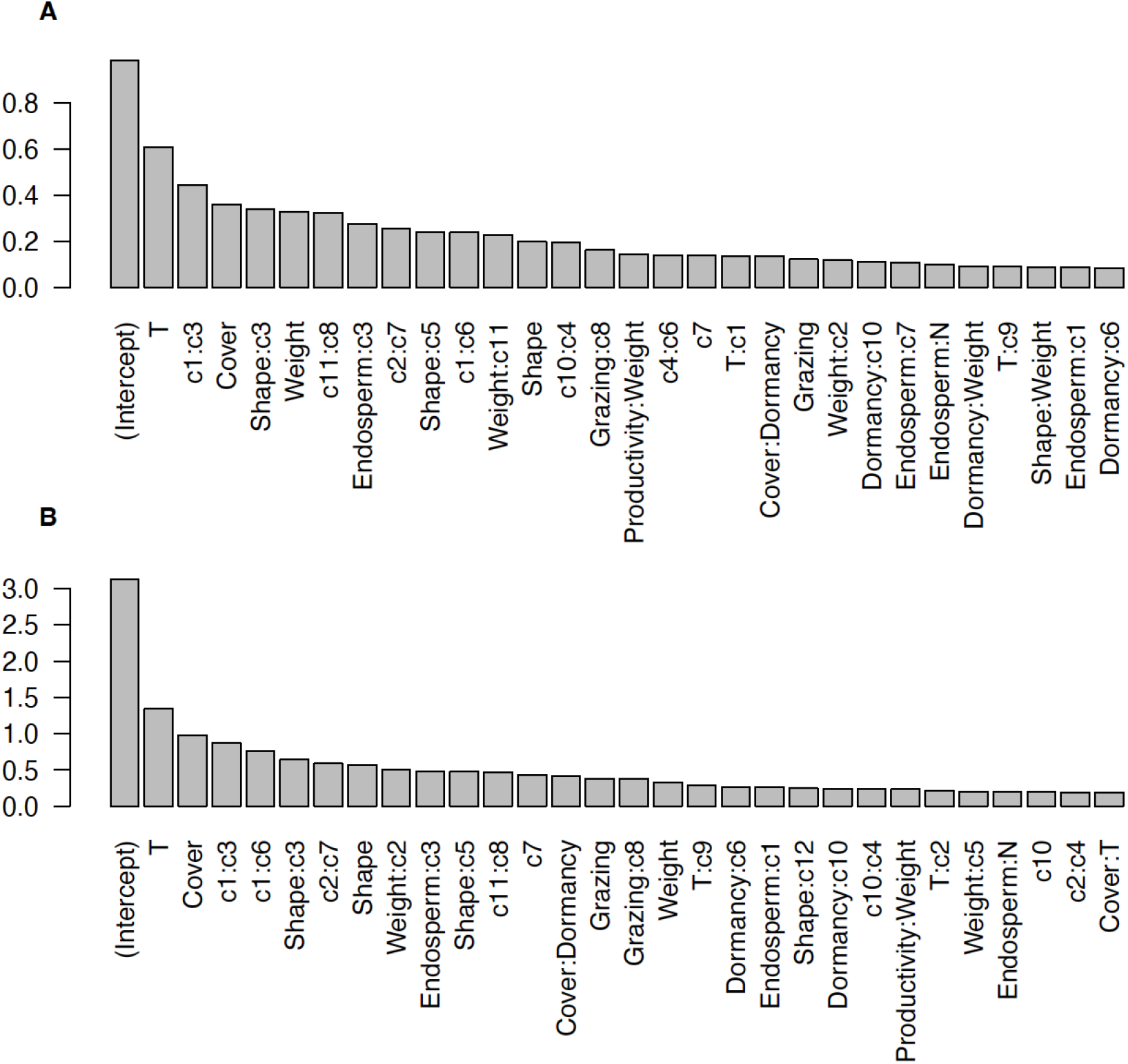

## Appendix S6.

Factors of Pearson correlation between predictors for SSB.

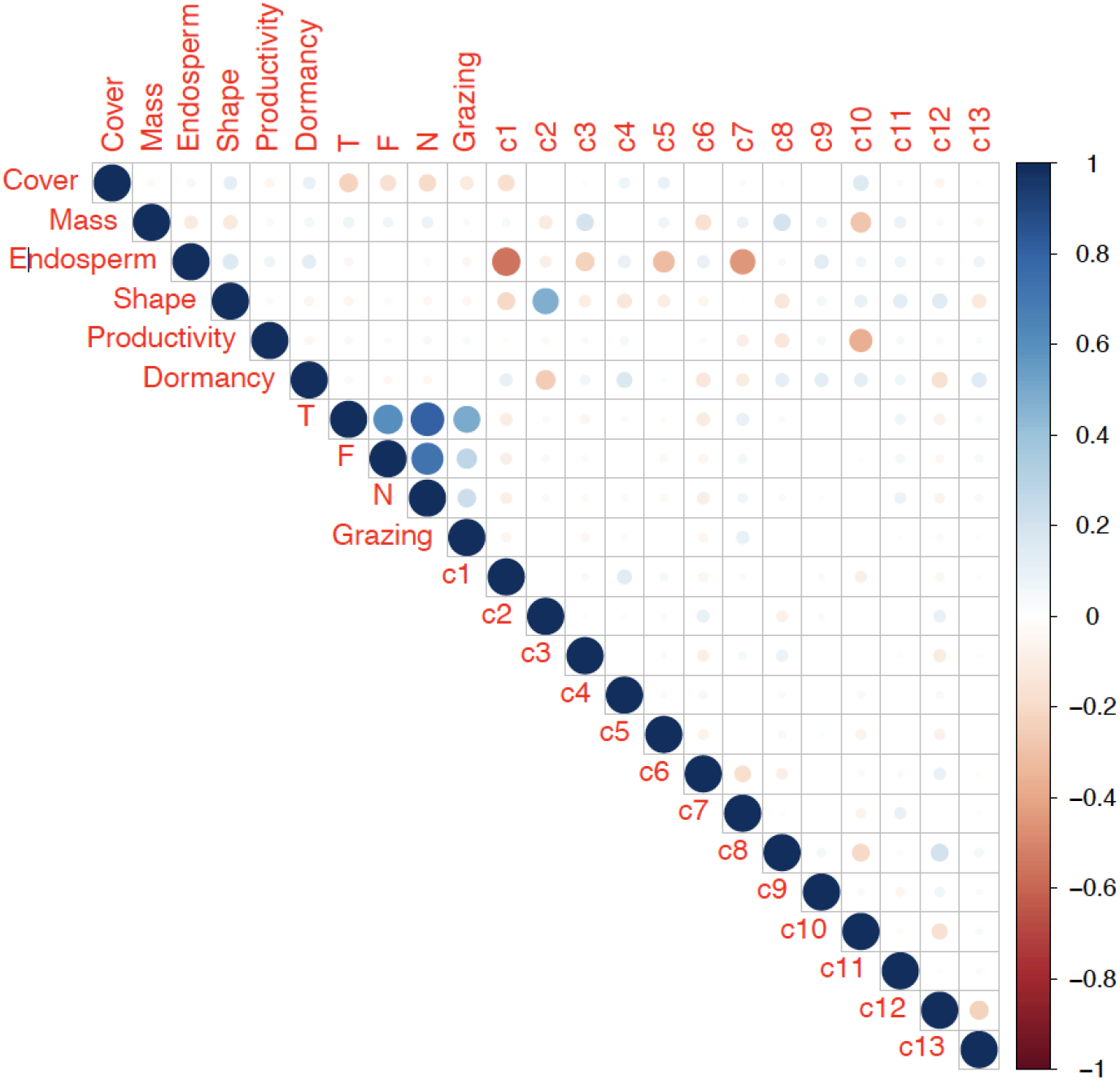

